# Comparative genomic analysis of Cluster AZ *Arthrobacter* phages

**DOI:** 10.1101/2025.05.26.655371

**Authors:** Amanda C. Freise, Kieran Furlong, Karen Klyczek, Andrea R. Beyer, Rebecca A. Chong, Nicholas P. Edgington, Bryan Gibb, Sarah J. Swerdlow, Madison G. Bendele, Isobel D. Cobb, Zachary Mitchell, Steve Cresawn, Amaya Garcia Costas, Adam D. Rudner

## Abstract

Bacteria in the *Arthrobacter* genus belong to the phylum Actinobacteria and are primarily soil-dwelling. Over 600 bacteriophages infecting *Arthrobacter* hosts have been isolated and sequenced, and genomic analyses show these phages to be highly diverse with mosaic genome architectures. We describe here a group of 32 *Arthrobacter* phages grouped in Cluster AZ, isolated on four different *Arthrobacter* strains all with siphoviral morphologies. The Cluster AZ phages exhibit a spectrum of diversity and can be subdivided into four subclusters. The diversity in minor tail protein and endolysin genes correlates partly with isolation host strain and may be predictive of the host range of these phages. Most of the Cluster AZ phages are temperate, form stable lysogens, and encode an integrase; however, an immunity repressor gene has not been identified. The intracluster diversity was analyzed in-depth at the whole genome level and through individual genes. As more *Arthrobacter* phages are isolated and analyzed they continue to provide new insights into phage evolution.

## Introduction

The soil-dwelling bacteria of the genus *Arthrobacter* belong to the Actinobacteria phylum. First defined as a group in 1928 and then officially proposed as a genus in 1947 (1), *Arthrobacter* spp. are obligate aerobes readily found in sewage and many types of soils including desert, agricultural, rhizosphere, Antarctic, and others (2,3). *Arthrobacter* spp. can metabolize a large diversity of organic and inorganic pollutants and have been of great interest in biotechnology and bioremediation applications, with numerous studies exploring *Arthrobacter* metabolic pathways and their ability to degrade a wide variety of pollutants such as pesticides and heavy metals (4,5). There are currently over 80 *Arthrobacter* species described with the type species being *A. globiformis* (6).

Like other bacteria, *Arthrobacter* spp. are infected by bacteriophages, and these phages likely play roles in the ecology, adaptations, and evolution of these soil microorganisms. Phages that can infect *Arthrobacter* hosts were isolated as early as the 1940s (7), although the first complete *Arthrobacter* phage genome sequence was not published until 2014 (8). Since then, hundreds of *Arthrobacter* phages have been isolated, mostly by students and faculty in the Science Education Alliance - Phage Hunters Advancing Genomics and Evolutionary Science (SEA-PHAGES) (9,10) and Phage Hunters Integrating Research and Education (PHIRE) (11) programs (12–14).

In 2017, Klyczek *et al*. published the first comparative genomics study of *Arthrobacter* phages, describing 46 phages isolated on *Arthrobacter sp.* (ATCC 21022) (15). These phages were organized into ten clusters and two singletons based on their average nucleotide identity (16). Clusters are groups of phages with closely related genomes (17–19), and some can be divided into subclusters based on genomic relationships; singletons are phages without any close relatives. The initial group of *Arthrobacter* phages shared very little nucleotide identity with phages isolated on other Actinobacteria hosts such as *Mycobacterium*, and the clusters were well separated genetically with relatively little gene sharing between clusters. Furthermore, almost all these phages appeared to be lytic, in contrast with other actinobacteriophages, where temperate phages are common (14). Over the past seven years, the number of *Arthrobacter* phage sequences has increased tenfold, with over 500 sequenced phages isolated on four *Arthrobacter* species (https://phagesdb.org). This larger collection of phages exhibits a higher level of diversity, comprising 30 clusters and four singletons, with several clusters predicted to have temperate life cycles.

One of the largest clusters of *Arthrobacter* phages is Cluster AZ. In an initial investigation, Kapinos *et al.* (2022) studied 12 AZ Cluster phages and reported a high degree of diversity within the cluster, along with shared gene content with *Microbacterium* phages in Clusters EH and EB (20). They also confirmed experimentally that some Cluster AZ phages were temperate and could generate stable lysogens. Here we report the comparative genomic analysis of an expanded set of 32 Cluster AZ phages, isolated from multiple geographically distinct locations in North America, which infect four different *Arthrobacter* hosts for isolation (21–26), revealing a continuum of diversity in the Cluster AZ phages.

## Results and Discussion

### Phage isolation and morphological characterization

Thirty-two phages were isolated on *Arthrobacter* bacterial hosts, of which twenty-seven phages were isolated on two strains of *A. globiformis*, three phages on *A. atrocyaneus,* and two phages on *A. sulfureus* (**Table 1**). All were isolated by plaque purification from soil samples collected across the United States and Canada (**Fig. 1A**) by students and faculty from 12 institutions participating in the SEA-PHAGES and PHIRE programs. Phages were isolated using either direct or enrichment isolation protocols (27). Plaque morphology was diverse among these phages, with some forming turbid plaques after a few days of incubation, consistent with temperate lifestyles (**Fig. 1B, C**). Phage particles visualized by transmission electron microscopy with negative staining showed that these phages have siphoviral morphology (**Fig. 1D**) as reported for other Cluster AZ phages (20), with isometric heads and non-contractile, flexible tails.

**Table 1.** Genometrics of Cluster AZ *Arthrobacter* phages.

| Phage Name | *Arthrobacter* Host Species | Subcluster | Length (bp)* | GC% | # Genes (tRNAs) | Plaque Morphology | Accession # |
| --- | --- | --- | --- | --- | --- | --- | --- |
| Adolin | *atrocyaneus* B-2883 | AZ1 | 43022 | 65.9 | 71 | Turbid | MN813676 |
| Adumb2043 | *globiformis* B-2880 | AZ1 | 43100 | 67.0 | 68 | Turbid | MT889375 |
| Amyev | *globiformis* B-2880 | AZ1 | 44284 | 67.9 | 68 | Turbid | OL549191 |
| Asa16 | *globiformis* B-2880 | AZ1 | 43601 | 66.6 | 69 | Turbid | MZ681506 |
| Crewmate | *globiformis* B-2880 | AZ1 | 44354 | 68.8 | 74 | Turbid | OL549189 |
| DrManhattan | *atrocyaneus* B-2883 | AZ1 | 42577 | 66.0 | 72 | Clear | MH834610 |
| DrSierra | *atrocyaneus* B-2883 | AZ1 | 42622 | 66.5 | 66 | Clear | MN908689 |
| Elezi | *globiformis* B-2880 | AZ1 | 43471 | 66.6 | 68 | Turbid | MT639653 |
| Eraser | *globiformis* B-2880 | AZ1 | 43608 | 66.0 | 69 | Clear | MZ747516 |
| Iter | *globiformis* B-2979 | AZ1 | 43963 | 67.4 | 70 | Turbid | ON208833 |
| Janeemi | *globiformis* B-2880 | AZ1 | 43877 | 67.1 | 69 | Turbid | ON970616 |
| Kaylissa | *globiformis* B-2880 | AZ1 | 44124 | 67.6 | 71 | Turbid | MZ005682 |
| KeAlii | *globiformis* B-2979 | AZ1 | 41850 | 65.5 | 67 (1) | Turbid | OK040777 |
| Lego | *globiformis* B-2880 | AZ1 | 43446 | 67.5 | 69 | Turbid | MT024869 |
| Lizalica | *globiformis* B-2880 | AZ1 | 43055 | 67.4 | 69 | Turbid | OL549192 |
| London | *globiformis* B-2880 | AZ1 | 43599 | 66.6 | 69 | Turbid | MT889366 |
| Niobe | *globiformis* B-2880 | AZ1 | 43602 | 66.7 | 69 | Turbid | MZ820087 |
| Phives | *globiformis* B-2880 | AZ1 | 44204 | 67.3 | 70 | Turbid | MT889376 |
| Powerpuff | *globiformis* B-2880 | AZ1 | 44651 | 67.6 | 71 | Turbid | MN703413 |
| Reedo | *globiformis*# | AZ1 | 41823 | 65.3 | 68 | Clear | OL455896 |
| Tbone | *globiformis*# | AZ1 | 44111 | 67.6 | 69 | Turbid | MW055910 |
| Tuck | *globiformis* B-2880 | AZ1 | 43992 | 67.0 | 69 | Turbid | OP820474 |
| VResidence | *globiformis* B-2880 | AZ1 | 42157 | 65.0 | 72 (2) | Turbid | OP434455 |
| Warda | *globiformis* B-2880 | AZ1 | 43792 | 67.6 | 70 | Turbid | OL549188 |
| YesChef | *globiformis* B-2880 | AZ1 | 43510 | 67.7 | 69 | Turbid | MT024871 |
| Yang | *globiformis* B-2979 | AZ1 | 43206 | 68.4 | 68 | Clear | MH834629 |
| Liebe | *globiformis* B-2979 | AZ2 | 45803 | 68.8 | 69 | Clear | MK061413 |
| Maureen | *globiformis* B-2979 | AZ2 | 45802 | 68.8 | 69 | Turbid | MH834619 |
| Snek | *globiformis* B-2979 | AZ3 | 40872 | 66.2 | 68 | Clear | PP208922 |
| Tweety19 | *globiformis* B-2979 | AZ3 | 40873 | 66.1 | 69 | Turbid | MT897906 |
| Emotion% | *sulfureus* ATCC 19098 | AZ4 | 45092 | 67.9 | 74 | Turbid | OQ709216 |
| VroomVroom% | *sulfureus* ATCC 19098 | AZ4 | 43230 | 66.9 | 67 | Turbid | OQ938592 |

\* Average genome length is 43,561 bp

**Figure 1.**
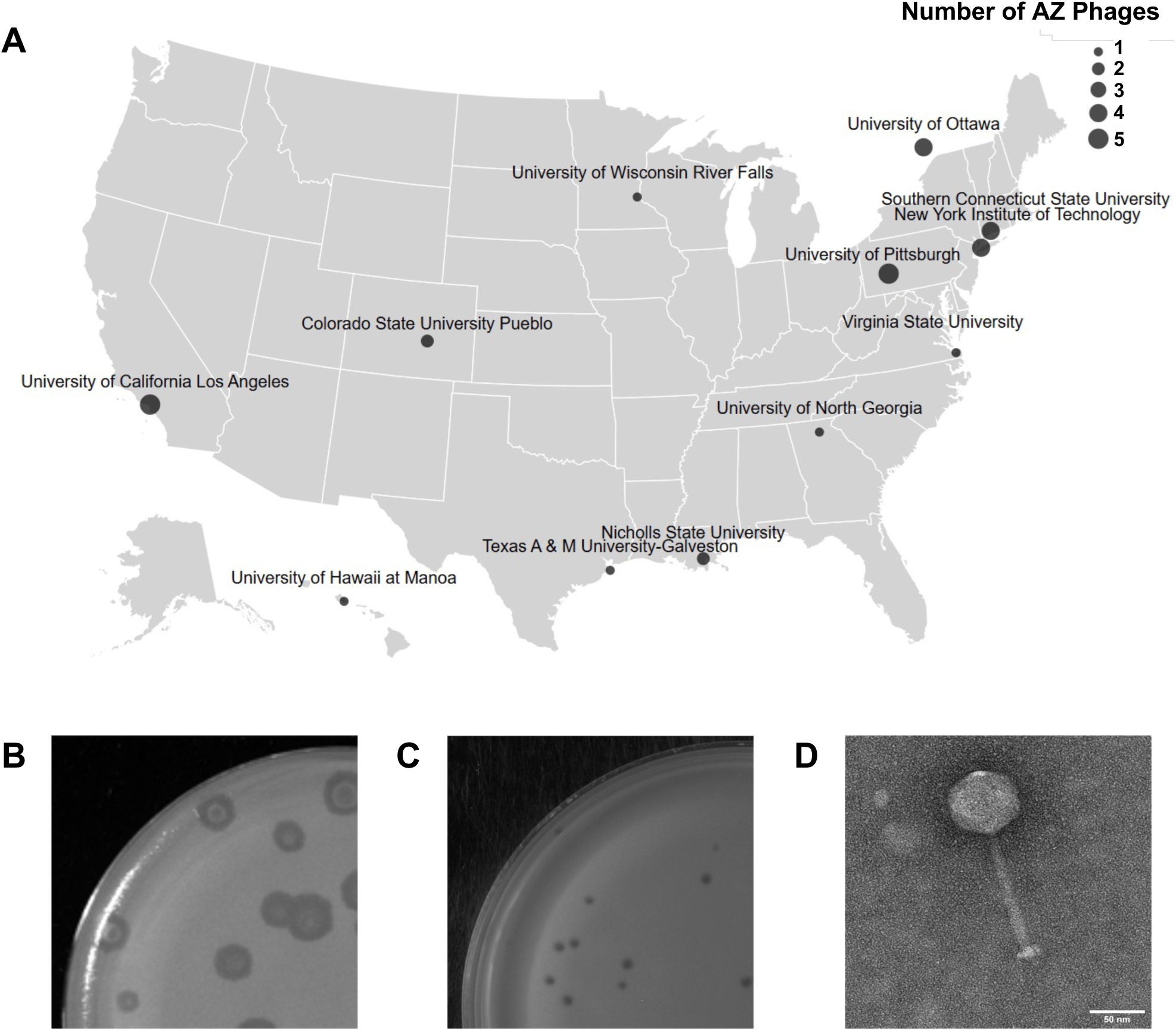
Geographical distribution of isolated Cluster AZ phages and phenotypic characterization. (A) Geographic distribution map of institutions that isolated Cluster AZ phages. (B, C) Variation in plaque morphologies of Cluster AZ phages. Some AZ phages form small clear plaques like DrManhattan (B) and others form turbid plaques like Lizalica (C). See **Table 1** for the plaque morphologies of all the AZ phages discussed in this paper. (D) Transmission electron micrograph of phage Crewmate with siphoviral morphology typical of Cluster AZ phages.

### Cluster AZ phage genometrics

The complete genomes of the 32 *Arthrobacter* phages were sequenced, and putative gene locations and functions were assigned based on bioinformatic analyses, as described previously (19,28,29). We harmonized the genome annotations performed by undergraduate and faculty participants in the SEA-PHAGES program with extant phage annotations for consistency in translation initiation codon and functional assignments. The process resulted in 89 changes in translation start site predictions, 61 changes in functional annotations, the addition of seventeen genes in gaps between previously annotated genes, and the assignment or adjustment of three programmed translational frameshifts in tail assembly chaperones. Intracluster annotation revision was facilitated by a custom Javascript notebook “Pharmonizer” (https://observablehq.com/@phage/pharmonizer) (**Fig. S1**) that indicates annotation discrepancies among related genes of the same ‘phamily’ or ‘pham’, where each pham contains genes with related amino acid sequences (18,30), and intragenic gaps where genes may have been overlooked. Pharmonizer can be broadly applied to add consistency to other intracluster genome harmonization.

Cluster AZ phage genomes vary in size, ranging from 40,872 bp (Snek) to 45,803 bp (Liebe), with an average genome size of 43,561 bp (**Table 1**). Lizalica is used as the primary benchmark for all genome comparisons discussed below (**Figures 1A and 2**). All genomes have defined ends with 10 or 11 base 3’ single-stranded DNA extensions. The %GC content also varies modestly, ranging from 65 - 68.8%, with an average %GC content of 67.1%. Genomes each have 65-75 predicted genes, with an average of 69 protein-coding genes per genome. With one or two exceptions per genome, most genes are transcribed rightwards (**Figure 2**). Two phages, KeAlii and VResidence, are predicted to contain tRNA genes; KeAlii has a tRNA-trp gene and VResidence has tRNA-trp and tRNA-Gln genes.

**Figure 2.**
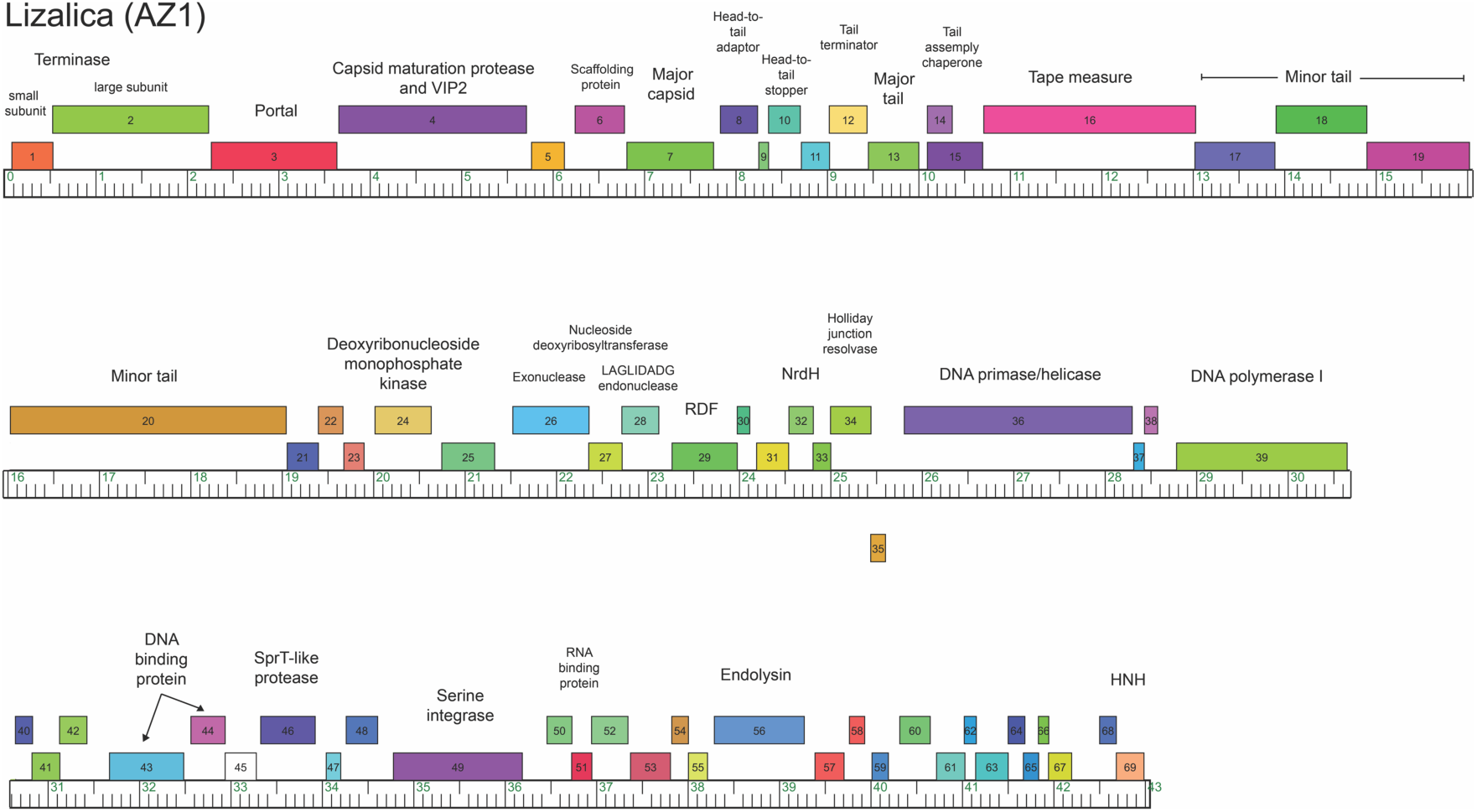
Lizalica genome map. Genome of phage Lizalica, a representative phage of the AZ Cluster. Genes are depicted as boxes and are colored according to their phamily designations (phams). White boxes (orphams) represent genes with no homologues present in the Actinobacteriophage database.

### Nucleotide and gene content comparisons

Phages are organized into clusters and subclusters based on proteomic equivalence quotient (PEQ) and Gene Content Similarity (GSC) (31) values calculated using PhamClust (17,32) that compare similarities between shared genes in each pairwise comparison of phages in the cluster. Pairwise gene content comparisons among all phages in Cluster AZ suggest these 32 phages subcluster into four groups (**Fig. 3A**, **Supplemental Table 1**). Within the largest subcluster, AZ1, GCS ranges from 62.8-99.3%. Subcluster AZ2 phages Liebe and Maureen share 100% GCS with each other, but only 52.9-63.3% GCS with phages in AZ1. Subcluster AZ4 phages Emotion and VroomVroom share 75.9% GCS with each other but <56% with phages in the other subclusters. The four subclusters vary in terms of size; AZ1 is much larger than AZ2, AZ3, and AZ4. In addition, the subclusters correlate only partially with the isolation host (**Table 1**). The larger Subcluster AZ1 includes phages isolated on two strains of *A. globiformis* and one strain of *A. atrocyaneus* (Adolin, DrManhattan, DrSierra), whereas the other subclusters are more distinct, with Subclusters AZ2 and AZ3 containing only *A. globiformis* phages, and Subcluster AZ4 including phages only isolated on *A. sulfureus*.

**Figure 3.**
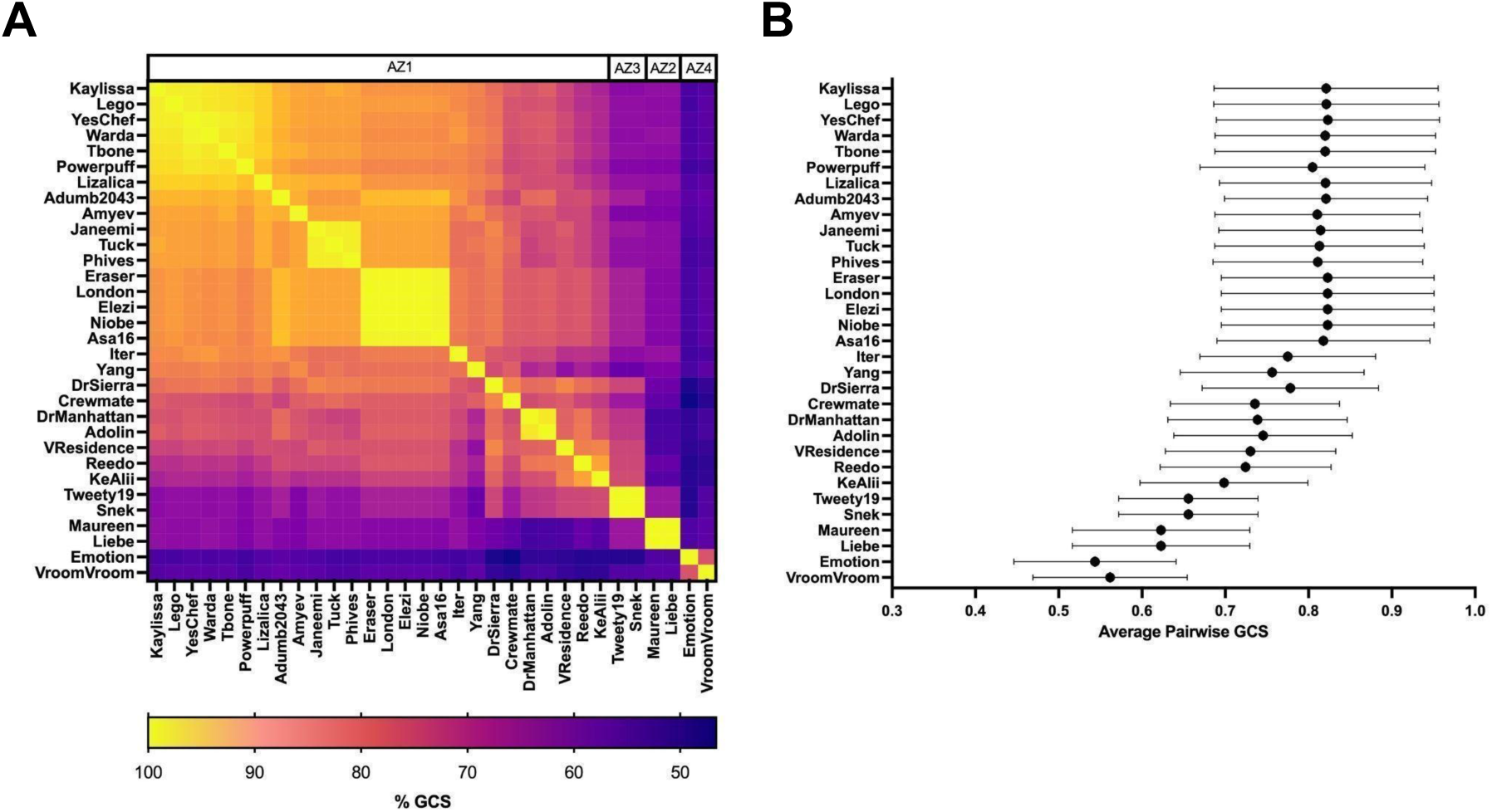
Gene content similarity in Cluster AZ. Pairwise gene content similarity (GCS) comparisons were performed for all Cluster AZ phages. (A) Heat map displaying pairwise GCS values. Phages are ordered according to subcluster designation. (B) Average pairwise GCS values for each phage are plotted. Error bars represent standard deviation. (n=31 comparisons per phage)

Some AZ1 phages are strikingly similar to each other. For example, Elezi, Asa16, Niobe, Eraser, and London all share at least 98.5% GCS. There is also a high level of GCS (90-98%) between Lizalica, Kaylissa, Lego, Powerpuff, YesChef, Tbone, and Warda. Other notable smaller sets of phages with high intragroup GCS include Adolin and DrManhattan (96.5%), which were both isolated on *A. atrocyaneus*. Outside of these instances, there are fewer discrete groupings of phages within AZ1 based on GCS. Approximately half of the phages (all in Subcluster AZ1) yielded average pairwise GCS values of 80% or greater (**Fig. 3B**). The rest of the phages yielded average pairwise GCS values that ranged from 54.2% (Emotion) to 80%. These patterns of similarity and dissimilarity in the GCS data illustrate a continuum of phage diversity as noted previously (19,31).

### Genome organization and pham content

The AZ genomes have a conserved gene organization in which the genes in the left arm code for structural proteins, while those in the right arm include genes that encode proteins needed for recombination, DNA metabolism, temperate life cycle, and many small genes without assigned functions (**Fig. 2 and 4**). The 32 genomes analyzed contain 2152 protein-coding genes grouped into 198 different phams (33) of which 49.6% have predicted functions. 25 phams are present in all 32 Cluster AZ genomes analyzed (**Fig. 4B**), including many with structural roles (e.g. Terminase, Portal protein, Major Capsid Protein) and DNA metabolism (e.g. DNA primase/helicase, nucleoside deoxyribosyltransferase). Most of these phams are widely distributed among Actinobacteriophages from other phage clusters (see below).

**Fig 4.**
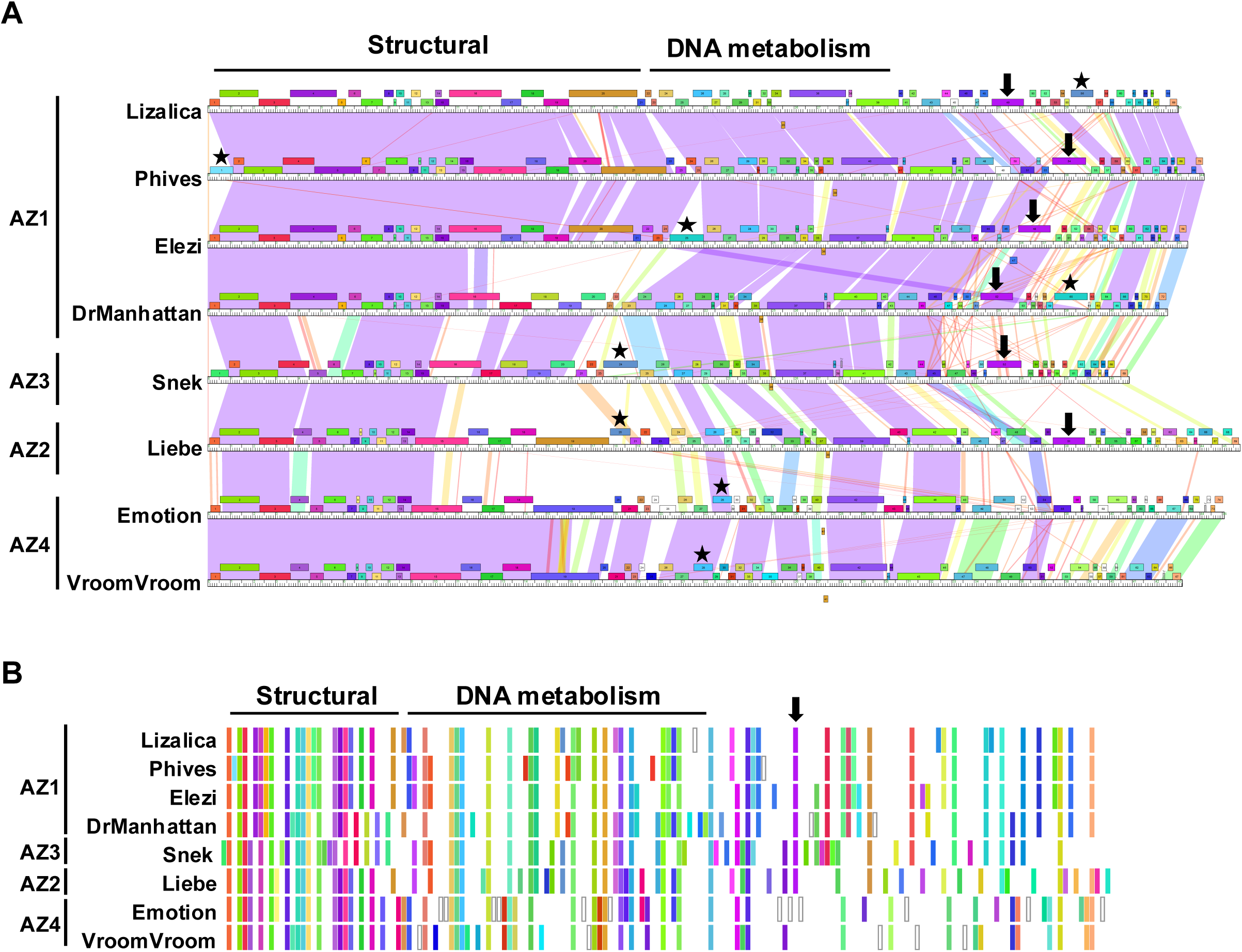
Genome maps and phamily matrix diagram of representative Cluster AZ phages. Genome maps of selected representative phages of each AZ subcluster. Maps were generated using the Phamerator.org (33) database Actinobacteriophage_4491 as described in Figure 2. Broad functional categories of each section of the genome are indicated at the top. Arrows indicate the position of Integrases and stars indicate the positions of Endolysins. (B) Phamily matrix diagram indicating mean pham positions in representative Cluster AZ phages. The diagram was generated using a custom Javascript notebook (Cresawn, manuscript in preparation; https://observablehq.com/@cresawn-labs/pham-matrix) that organizes and aligns phams based on their mean position across the selected genomes (database Actinobacteriophage_4491) and illustrates pham conservation across the selected genomes. Phams are depicted as colored bars corresponding to their respective pham as in the Phamerator map in (A). As in (A) arrows indicate the position of Integrases. The endolysins are not marked because of their varied genomic position.

### Relationships to other Actinobacteriophages

To determine the relationship between AZ Cluster phages and other clusters of *Arthrobacter* phages we created a gene content network phylogeny that assesses relationships based on shared phams of the AZ Cluster phages (**Fig. 5**). We included two clusters of *Microbacterium* phages, EB and EH, that are closely related to cluster AZ phages (20). The Cluster AZ phages form a distinct group among the *Arthrobacter* phages, with phages in this cluster only sharing substantial inter-cluster gene content with *Arthrobacter* Cluster FP phages. In contrast, Clusters AY, FA, FB, FF, FN, FO, and Singleton TripleJ, share substantial inter-cluster gene content as indicated by the interconnecting branches among these clusters. This pattern is consistent with the observation that while Cluster AZ phages share 24-30**%** gene content similarity (GCS) with *Arthrobacter* phages in Cluster FP, they are more closely related to *Microbacterium* Cluster EH phages (35.2-37.3% GCS) (20).

**Fig 5.**
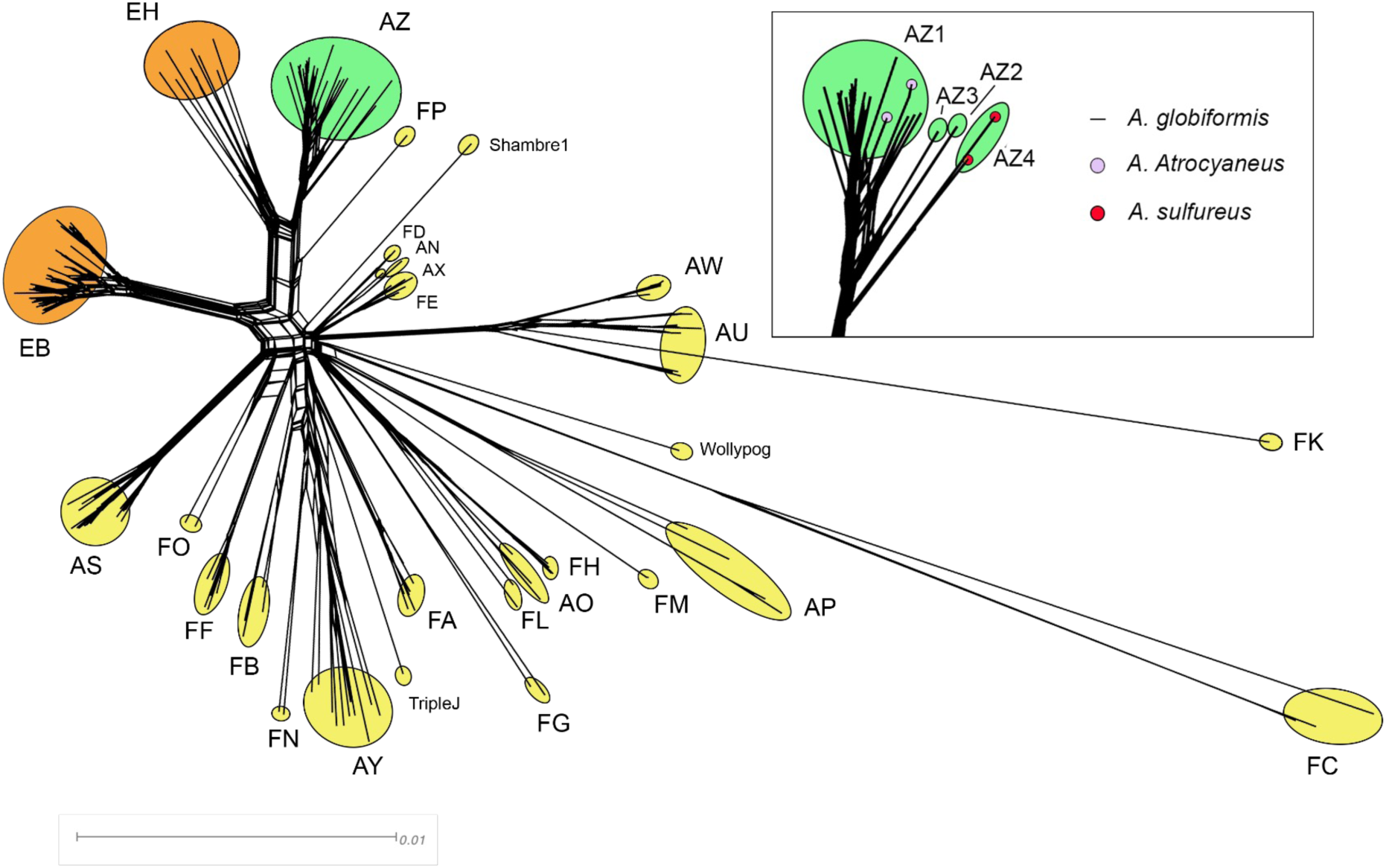
A network phylogeny of selected *Arthrobacter* and *Microbacterium* phages. The predicted proteins of all phages in the database Actinobacteriophage_4491 were sorted into phams according to shared amino acid sequence similarities using PhaMMseqs (30,33). (A) Each genome of the phages isolated on *A. globiformis, A. atrocyaneous, and A. sulfureus*, and phages in *Microbacterium* Clusters EB and EH, was then assigned values reflecting the presence or absence of each pham member. The genomes were compared and displayed using Splitstree (67). The clusters are indicated with colored ovals (green, Cluster AZ; yellow, other *Arthrobacter* phages; orange, *Microbacterium* phages) with singleton phages specified by name. The scale bar indicates 0.01 substitutions/site. (B) The inset shows subcluster designations for Cluster AZ phages. Small colored circles at the tips indicate the phages isolated on *A. atrocyaneus* and *A. sulfureus,* as noted in the key. All other phages were isolated on *A. globiformis*.

Two Cluster FP phages, BaileyBlu and CallinAllBarbz, were compared with Clusters AZ, EH, and FP phages (**Fig. 6**). The left arm of the genomes of the Cluster FP phages have little or no nucleotide sequence similarity with Cluster AZ phages, and the shared genes are primarily among the non-structural genes in the right halves of the genomes. In contrast, Cluster EH genomes share nucleotide similarity with Cluster AZ phages in this region, which primarily includes genes encoding structural proteins. Both Cluster FP and EH phages are similar to Cluster AZ phages in the region of the genome encoding DNA metabolism proteins. The genes having the highest sequence similarity between the AZ and FP phages include Cas4 family exonuclease (BaileyBlu *25*) recombination directionality factor (BaileyBlu *30*), DNA primase/helicase (BaileyBlu *38*), and DNA polymerase I (BaileyBlu *41*) (**Fig. S2)**. BaileyBlu also includes two tRNA genes, a Trp and a Met, near the end of the right arm of the genome, similar to the AZ phage VResidence which has Trp and Gln tRNAs in the same region.

**Figure 6.**
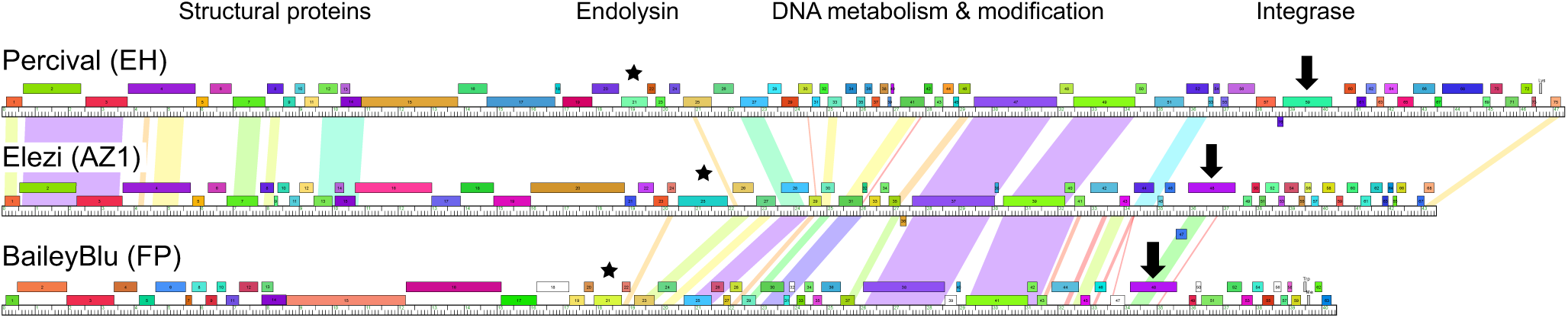
Pairwise alignment of Cluster EH, AZ, and FP phage genomes. Representative genomes from *Microbacterium* Cluster EH and *Arthrobacter* Clusters AZ and FP are shown, using Phamerator.org (33) database Actinobacteriophage_4491. Pairwise nucleotide sequence similarities are shown by spectrum coloring between the genomes (violet indicates the highest similarity, red the least similarity). Genes are depicted as boxes and are colored according to their phamily designations. White boxes represent genes with no close relatives in the database (orphams). Broad functional categories of each section of the genome are indicated at the top. The endolysin genes are indicated with asterisks, and the integrase genes are shown by arrows.

### Genes for virion structure and assembly

Genes encoding structural proteins are located in the left arm of the Cluster AZ genomes and include small and large terminase subunits, portal protein, major capsid protein, head-to-tail adaptor and stopper, tail terminator, major tail protein, tail assembly chaperones, tape measure protein, and minor tail proteins. The presence of these functions, in this gene order, is typical for siphoviral bacteriophages. In Subcluster AZ1 phages, the capsid maturation protease is fused with a VIP2-like ADP ribosyltranserase toxin (Lizalica *4*), while in AZ2, AZ3, and AZ4 phages only the protease domain is present (Liebe *4*). The fused gene is shared with Cluster EH phages (20,34). The role of this protein in phage growth is not known. The AZ phages have a terminase-associated HNH endonuclease, found at the right end of the genome and is joined with the terminase, located at the left end of the genome, after genome injection and circularization during the initial stage of phage infection (35).

Capsid genes are highly conserved in the AZ cluster and in many other clusters of phages infecting multiple host genera, whereas proteins associated with tail structures diverge between the four AZ Subclusters and are shared with fewer other clusters. For example, the portal (Lizalica *3*) pham is shared with phages that infect *Mycobacterium*, *Gordonia*, *Streptomyces*, and *Rhodococcu*s. The wide distribution of these conserved, essential genes suggests a shared evolutionary history for these Actinobacteriophages. The major capsid protein pham (Lizalica *7*) has a similar cluster distribution but has fewer members and is found in a smaller but still diverse set of clusters (BD, BQ, DU, L, EH and M). In contrast, the tape measure protein (Lizalica *16*) has a narrower cluster distribution, only found in Cluster AZ and *Microbacterium* Cluster EJ phages. None of the structural protein phams are unique to Cluster AZ.

### Minor Tail Proteins

All Cluster AZ phages have a group of four minor tail proteins (MTPs) (**Fig. 7A**; Lizalica *17-20*, **Fig. 2 and 4**). The minor tail proteins encode the phage’s tail tip structural components, which is essential for host recognition and triggering DNA injection into the host (36). Since the Cluster AZ phages were isolated on four different hosts (**Table 1**) we examined if any of the four MTP genes correlated with the isolation host. Dendrograms based on multiple sequence alignment of the four MTP genes (**Fig. 7B-E**) show the first three MTPs (i.e Amyev *17, 18, 19*) are shared between the second and third gene structure groups (**Fig. 7A**), while the fourth MTP (i.e Amyev *20*) is distinct between the three gene structure groups, suggesting that the fourth MTP may be involved in host selection. Phages with the second gene structure infect *A. globiformis* (NRRL B-2979 and B-2880 strains) and those with the third gene structure only infect *A. sulfureus*, supporting the model that the fourth MTP may contribute to host specificity. Further, the fourth MTP in the first gene structure group is also unique.

**Figure 7.**
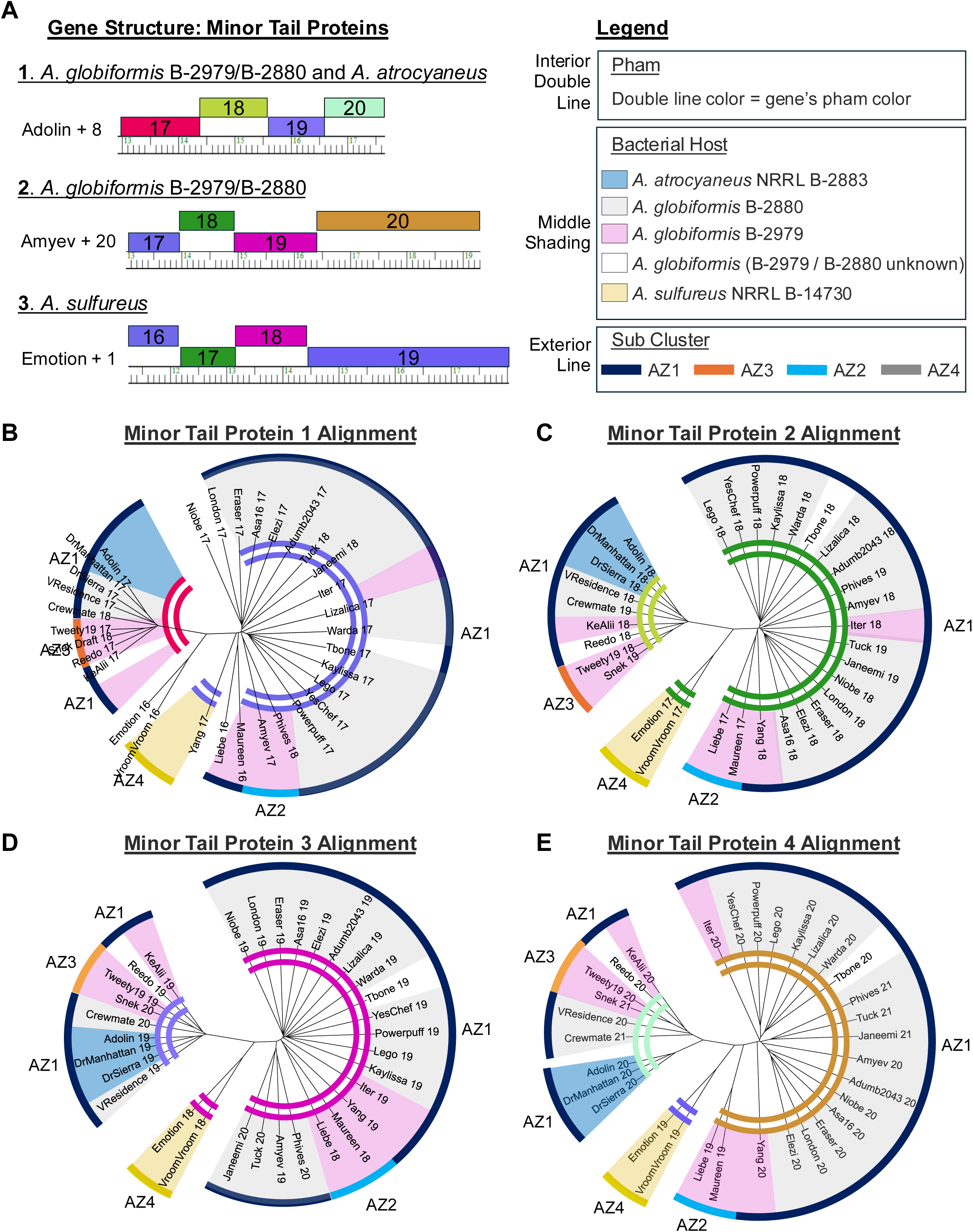
Phamerator map and multiple sequence alignments (MSAs) of the four minor tail proteins in AZ cluster phages. (A) Phamerator map showing the three different structures of the four minor tail genes in the AZ cluster, colored by pham. (B-E) MSA of the minor tail proteins. Solid exterior ring indicates the AZ subcluster, the middle-shaded ring indicates the phage isolation host and the interior double ring is colored to match its pham as in (A).

The grouping of the AZ1 and AZ3 phages is intriguing. Despite being isolated on three different hosts (*A. globiformis* NRRL B-2880, *A. globiformis* NRRL B-2979 and *A. atrocyaneus* NRRL B-2883), they share similar MTP gene structures (**Fig. 7A**) and cluster together in each of the four MTP alignments (**Fig. 7B-E**). We hypothesize that these phages may have a broader host range, and specifically, phages isolated on *A. globiformis* NRRL B-2880 could potentially infect either *A. globiformis* NRRL B-2979 or *A. atrocyaneus*, given their MTP similarity.

Most *A. globiformis* NRRL B-2979 phages group together. However, Iter is an exception. Either it clusters among the *A. globiformis* NRRL B-2880 phages, as observed in the first and second MTP alignments (**Fig. 7B-C**) or it borders them, as seen in the third and fourth MTP alignments (**Fig. 7D-E**). Iter’s unique position in the MTP alignments may indicate an evolutionary bridge between phages which infect *A. globiformis* NRRL B-2979 and *A. globiformis* NRRL B-2880.

Cluster FP phages, isolated on *A. sulfureus* ATCC 19098, share no MTPs with Cluster AZ phages, including the Subcluster AZ4 phages isolated on the same host. This suggests that the fourth MTP in cluster AZ4 phages is not a strict requirement for infection of *A. sulfureus* and highlights the complexity of host specificity. The MTPs in Cluster EH phages, isolated on *M. foliorum*, are also distinct from those in both Cluster AZ and FP phages.

### Lysis functions

Phage endolysins are required to disrupt the bacteria cell wall during cell lysis and release of new phages. These enzymes degrade the peptidoglycan layer by hydrolyzing amide or peptide linkages. *Arthrobacter* phage endolysins are diverse and modular (15) as described for the lysin A proteins in *Mycobacterium and Gordonia* phages (37,38). Many have three domains, typically including a peptidoglycan hydrolase/muramidase, an amidase, and a peptidoglycan binding domain, while others have only subsets of these. In some phages, these domain functions are encoded by separate, adjacent genes.

There are five types of predicted endolysins encoded in the Cluster AZ phage genomes, each with a distinct domain organization (**Fig. 8A, Supplemental Table 2**). These endolysins are found in three distinct genome locations **(Fig. 4A**). The endolysin types and locations partially correlate with subcluster assignment and isolation host.

**Fig 8.**
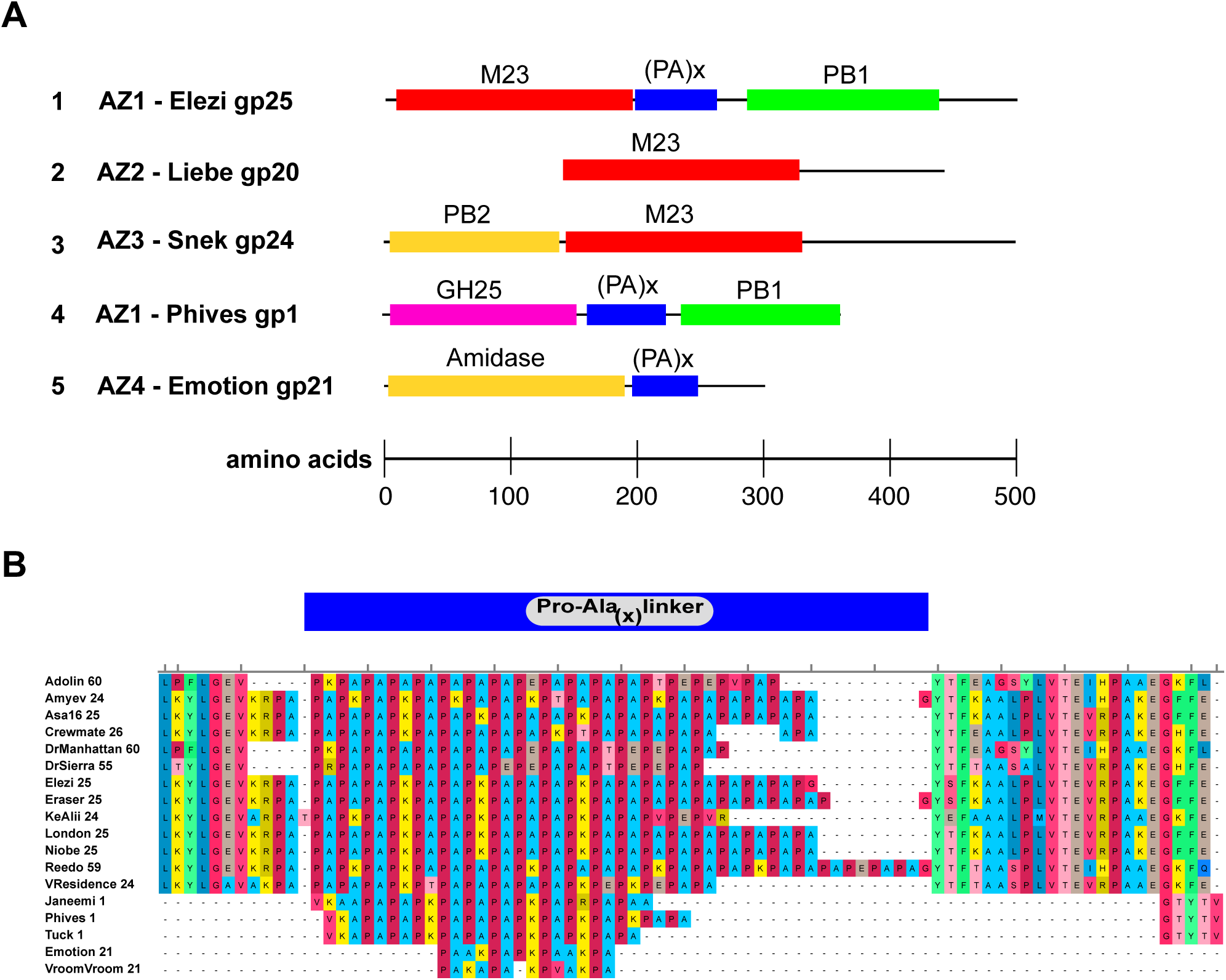
Five distinct types of endolysins in Cluster AZ phages. (A) Domains and organization of the five different endolysins observed in Cluster AZ phages. Domain abbreviations: M23: M23 peptidase-like domain; (PA)x: proline-alanine repeat; PB1: peptidoglycan binding domain type 1; PB2: peptidoglycan binding domain type 2; GH25: GH25-like muramidase domain. The scale indicates the amino acid number. (B) Multiple sequence alignment of the central region of AZ phage endolysins, showing the proline-alanine repeat region. Alignments were generated using MUSCLE.

One type of endolysin, illustrated by Elezi gp25, is found in 13 Subcluster AZ1 phages. This protein has an N-terminal domain similar to peptidase M23 and a C-terminal peptidoglycan-binding domain. These domains are separated by a central region containing several proline-alanine repeats (**Fig. 8B**). In nine of these phages, this endolysin gene is located in a canonical position, just downstream of the minor tail protein genes in the center of the genome (**Fig. 4A**).

However, in the other four phages, this gene is located in the right arm (e.g. DrManhattan *60*). Three of the four phages with the alternate endolysin location (Adolin, DrManhattan, DrSierra) were isolated on *A. atrocyaneus*, while the fourth, Reedo, was isolated on *A. globiformis* NRRL B-2880.

A second type of endolysin is found in the other Subcluster AZ1 phages and the Subcluster AZ2 phages. These proteins, represented by Liebe gp20, contain only one functional domain identified by HHPred, with strong homology to peptidase M23 and similar to the N-terminal peptidase domain in Elezi gp25 (**Fig. 8A**). The C-terminal region likely contains a novel functional domain with no homologues in the databases. In the Subcluster AZ2 phages, this endolysin gene is in the central location near the tail genes, while in the AZ1 phages, it is located in the right arm, similar to DrManhattan *60* (**Fig. 4A**).

The endolysin in AZ3 phages represents a third type of endolysin. Snek gp24 contains an M23 peptidase-like domain similar to that in Liebe gp20 and also has an N-terminal amidase domain that may include a peptidoglycan binding region (**Fig. 8A**).

The predicted endolysin in Phives, Janeemi, and Tuck, all Subcluster AZ1 phages, is distinct from the others in Cluster AZ, both in domain type and location. It is the first gene in these genomes and encodes a protein (represented by Phives gp1) with an N-terminal muramidase domain and C-terminal peptidoglycan binding domain (**Fig. 8A**). This pham is shared with other Arthrobacter phages in Cluster FG, as well as *Microbacterium* phages in Cluster EA3.

The phages isolated on *A. sulfureus*, in Subcluster AZ4, also contain a distinct type of endolysin located in the central genome region. Emotion gp21 has one identifiable domain, N-acetylmuramoyl-L-alanine amidase, in the N-terminus of the protein. Like Elezi gp25, there may be a novel functional domain in the C-terminal portion (**Fig. 8A**).

The endolysins encoded by Cluster AZ phage genomes are distinct from those found in related phages. The predicted endolysin gene in Cluster FP phages (BaileyBlu *21*) is only found in that cluster and has no other homologues in the database (**Fig. 6**). This endolysin has an N-terminal peptidase domain similar to M15 and a C-terminal peptidoglycan-binding domain.

Cluster EH phage genomes encode an endolysin (Percival *21*) found in several other *Microbacterium* phage clusters but not in any *Arthrobacter* phage genomes (**Fig. S2**). This Cluster EH endolysin has one identifiable domain, a carboxypeptidase in the N-terminal half of the protein. In both FP and EH phages, the endolysin genes are in the canonical location, just downstream of the minor tail protein genes (**Fig. 6, Fig. S2**).

Cluster AZ phages do not contain genes encoding homologues of lysin B, consistent with other *Arthrobacter* as well as *Microbacterium* phages (15,34). These hosts do not have the mycolic acid-rich outer membrane, found in *Mycobacterium* and *Gordonia* hosts, which requires lysin B for cell lysis (37).

Effective cell lysis also requires a holin that allows access to the cell wall through the plasma membrane. We cannot identify specific holin genes in these phages, but there are two or three well-conserved genes in each phage encoding proteins with multiple transmembrane helices, typical for holin proteins, that may serve this function (e.g. Elezi *22* - *24*). As in many phages (38,39), these potential holins are located adjacent to endolysin genes. However, in phages with downstream endolysin genes (e.g. Lizalica *56*) these putative holin genes are still located downstream of the minor tail genes, complicating a definitive annotation of these genes as holins.

### DNA metabolism functions

Several genes encoding proteins predicted to be involved in DNA biosynthesis and metabolism are highly conserved across all Cluster AZ phages in the right arm of the genome. These functions include DNA replication enzymes such as primase/helicase (Lizalica *36*) and DNA polymerase I (Lizalica *39*), as well as nucleoside metabolism enzymes deoxynucleoside monophosphate kinase (Lizalica *24*) and nucleoside deoxyribosyltranserase (Lizalica *27*).

Additional conserved genes encode a Cas4 family exonuclease (Lizalica *26*), LADLIDADG family endonuclease (Lizalica *28*), NrdH-like glutaredoxin (Lizalica *32*), Holliday junction resolvase (Lizalica *34*), and an HNH homing endonuclease as the last gene (Lizalica *69)*. These gene phamilies are also shared with phages in Clusters FP and EH (20), reflecting the shared evolutionary relationships among these clusters.

Other genes are less well conserved and illustrate the higher level of diversity in this region of the genome. These genes include a DNA ligase (Lizalica *41*) that is present in the genomes of most AZ1 phages (except Iter, KeAlii, VResidence) but not in other AZ Subclusters. An RNA binding protein (Lizalica *51*) is found in all Subcluster AZ1, AZ2, and AZ3 genomes, as well as in Cluster EH and FP, but not in Subcluster AZ4. A second gene encoding an RNA-binding protein is located just downstream (Adolin *56*) and is present in several AZ1 and both AZ3 phages, but not in AZ2 or AZ4. Subcluster AZ4 phage VroomVroom encodes a ThyX thymidylate synthase (gene *35*) that is not found in any other AZ phage genome but is present in several other phage clusters isolated on a range of host genera.

### Lysogeny

Some Cluster AZ phages such as Lego, Powerpuff, and YesChef are temperate and form stable lysogens (20). They encode a serine integrase for chromosomal integration but an immunity repressor gene has not been identified. The integrase is highly conserved among the AZ1, AZ2, and AZ3 phages, and the genes are located in the right arm of the genome (Lizalica *49*). All AZ phages, including AZ4, also encode a recombination directionality factor (Lizalica *29*) that is required for the excision of phage DNA (40). Subcluster AZ4 phages Emotion and VroomVroom do not have an identifiable integrase gene and may not be able to integrate into the host genome. However, both phages form turbid plaques, suggesting that they may form lysogens without genome integration (41,42) or using an unidentified integrase (43). We have also isolated lysogens of several additional AZ1 phages (Rudner, manuscript in preparation), suggesting that many AZ phages can form stable lysogens.

Cluster AZ phages contain four predicted DNA binding proteins that are candidates for functioning as immunity repressors. One, Lizalica *43*, is conserved in all AZ cluster members, and is present as a tandem duplication in AZ4 phages VroomVroom and Emotion (**Fig. 9A**).

**Fig. 9.**
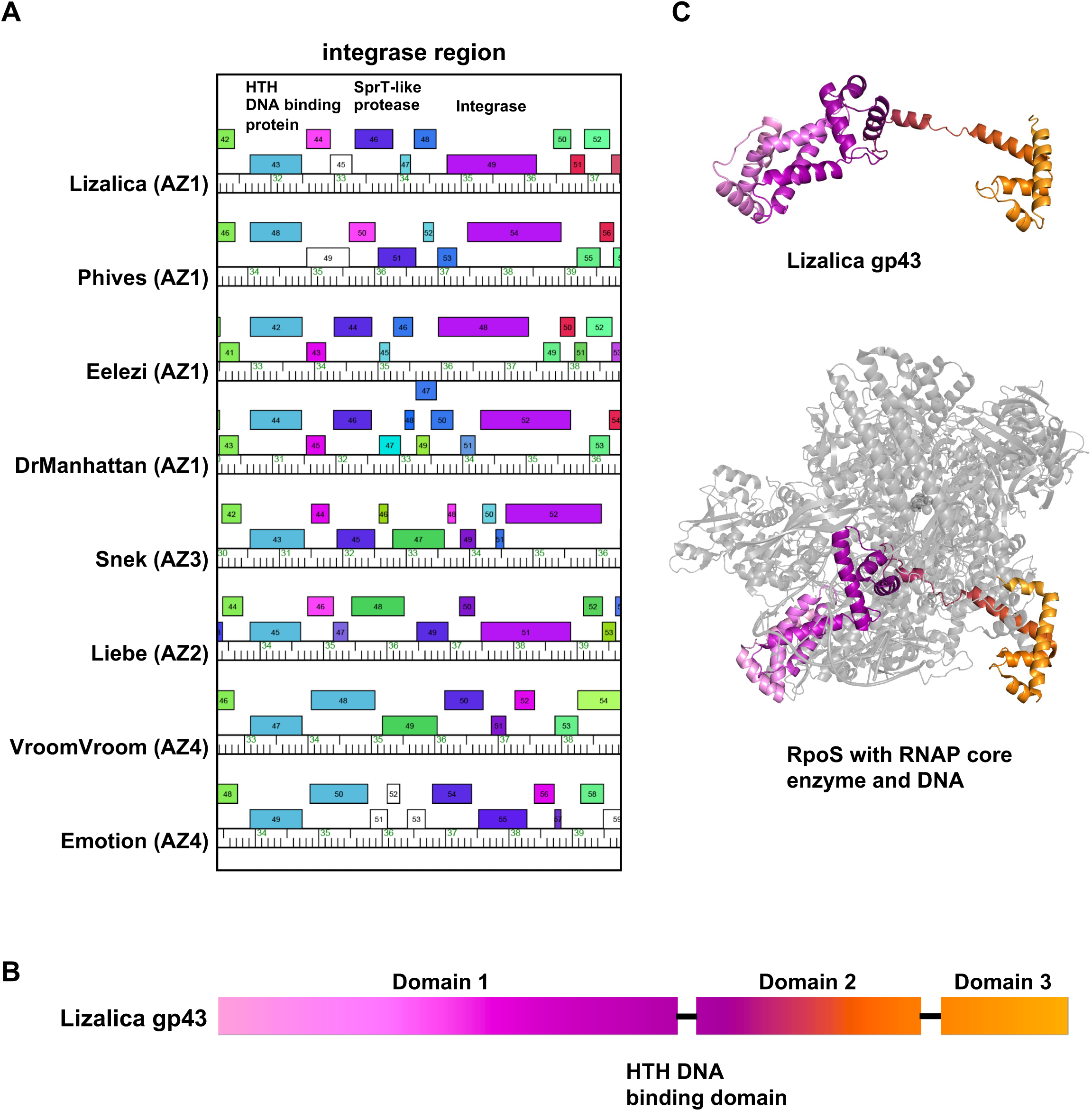
Conserved genes upstream of the Integrase in AZ phages. (A) Partial genome maps showing regions between HTH DNA binding proteins (Lizalica *43*) and serine integrases (Lizalica *49*) in representative phages. (B) Domain organization of the HTH DNA binding protein. A helix-turn-helix motif between domains 1 and 2 forms the predicted DNA binding domain in Lizalica gp43 which is homologous to the HTH domain of sigma factors. The color-coding matches that in the predicted and experimental structures in (C). (C) Comparison of the Alphafold3 predicted structures of Lizalica gp43 and the crystal structure of RpoS within the RNAP holoenzyme (5IPL (chain F)).

Lizalica gp43 has significant similarity to sigma factors (HHpred >99% probability) and a predicted Alphafold3 structure reveals three structural domains with a HTH DNA binding domain formed between domains 1 and 2 (**Fig. 9B and C**). These domains are present in the crystal structure of the *E. coli* RpoS sigma factor, the top match for Lizalica gp43 in the PDB100 database using the Foldseek server (**Fig. 9C)** (44), and the predicted structure closely matches the structure of RpoS bound to the RNAP holoenzyme, suggesting that Lizalica gp43 may function by interacting with RNA polymerase.

A second DNA binding protein, Lizalica *46,* is also conserved in all AZ cluster members and is homologous to a SprT (Spartan) protease. SprT proteases contain a C-terminal DNA binding domain and, in eukaryotes, are recruited to sites of protein-DNA crosslinks where their proteolytic activity is needed to degrade the crosslinked protein and repair these lesions (45).

Lizalica gp46 contains the conserved HEXXH metal binding motif found in SprT proteases and its predicted Alphafold3 structure reveals similarity to a *Pyrococcus abyssi* metalloprotease (tPDB 4JIU; top match in the PDB100 database using the Foldseek server) and the mammalian SprT protease domain (HHpred >99% probability). The small C-terminal DNA binding domain of Lizalica gp46 matches a variety of nucleic acid interacting proteins and, like the human SprT protease, the predicted DNA binding domain is formed from a short helical segment positioned below a two stranded **β**-sheet. These data support the assigned function of a SprT-like protease and highlight its resemblance in structure, and we predict in function, with its well-characterized mammalian homolog.

The AZ cluster Spr-T-like protease is ∼198 amino acids in all AZ phages examined, except for phage Liebe which contains a one base insertion in the 5’ end of the gene such that a downstream start site causes a 33 amino acid N-terminal truncation (**Fig. S3**). Phages Liebe and Maureen were found in the same soil sample and differ only by this insertion and three additional nucleotide changes (in Liebe *19*, *40* and *55;* **Fig. S3**). Notably, Liebe forms clear plaques (https://phagesdb.org/phages/Liebe/), while Maureen forms turbid plaques (https://phagesdb.org/phages/Maureen/), supporting a model in which the N-terminal protease domain of Lizalica gp46 (and Maureen gp49) may be required for efficient lysogeny.

The other two predicted DNA binding proteins with no defined functions in DNA metabolism are not present in all AZ cluster phages, though one, Lizalica *44* (present in Lizalica, Phives and Snek) (**Fig. 2**, **Fig. 9A**) is located just downstream of the conserved HTH DNA binding protein. This protein is smaller than Lizalica *43* and, although it is predicted to also contain an HTH DNA binding motif (HHpred 98-99% probability), it lacks structural similarity outside the DNA binding domain with other classes of transcriptional regulators.

In most mycobacterial phages, the repressor and integrase genes are in the same genomic region (46,47), so the position of two conserved DNA binding proteins, Lizalica *43* and *46*, upstream of the integrase, Lizalica *49*, may suggest that this region plays a role in regulating lysogeny. Other than Lizalica *43*, *46* and *49*, the genetic composition in this region is highly variable and contains several repeat sequences (data not shown) that might be indicative of the mobility of this cassette and its evolutionary history. Homologues of Lizalica *43* are found in EB, EH and FP cluster phages. EB and EH cluster phages share the AZ cluster genomic organization of this region, while FP phages lack the SprT-like protease.

### Quantitative Pham Analysis

To better understand gene mosaicism and evolutionary history in the genomes of AZ cluster phages, we evaluated the distribution of protein-coding genes within the AZ cluster. Our analysis revealed that Cluster AZ phage genomes contain 198 different phams, of which 105 are only found AZ cluster phage genomes (**Fig. 10A**) and these unique phams all have no known function. 41% of these are orphams, genes found only in an individual genome, with no homologues in the Actinobacteriophage database. Most of these genes are small (<500bp) and are located in the right arm of the genome. Cluster-specific phams may reflect novel adaptations in these phages to their hosts and environment.

**Fig. 10.**
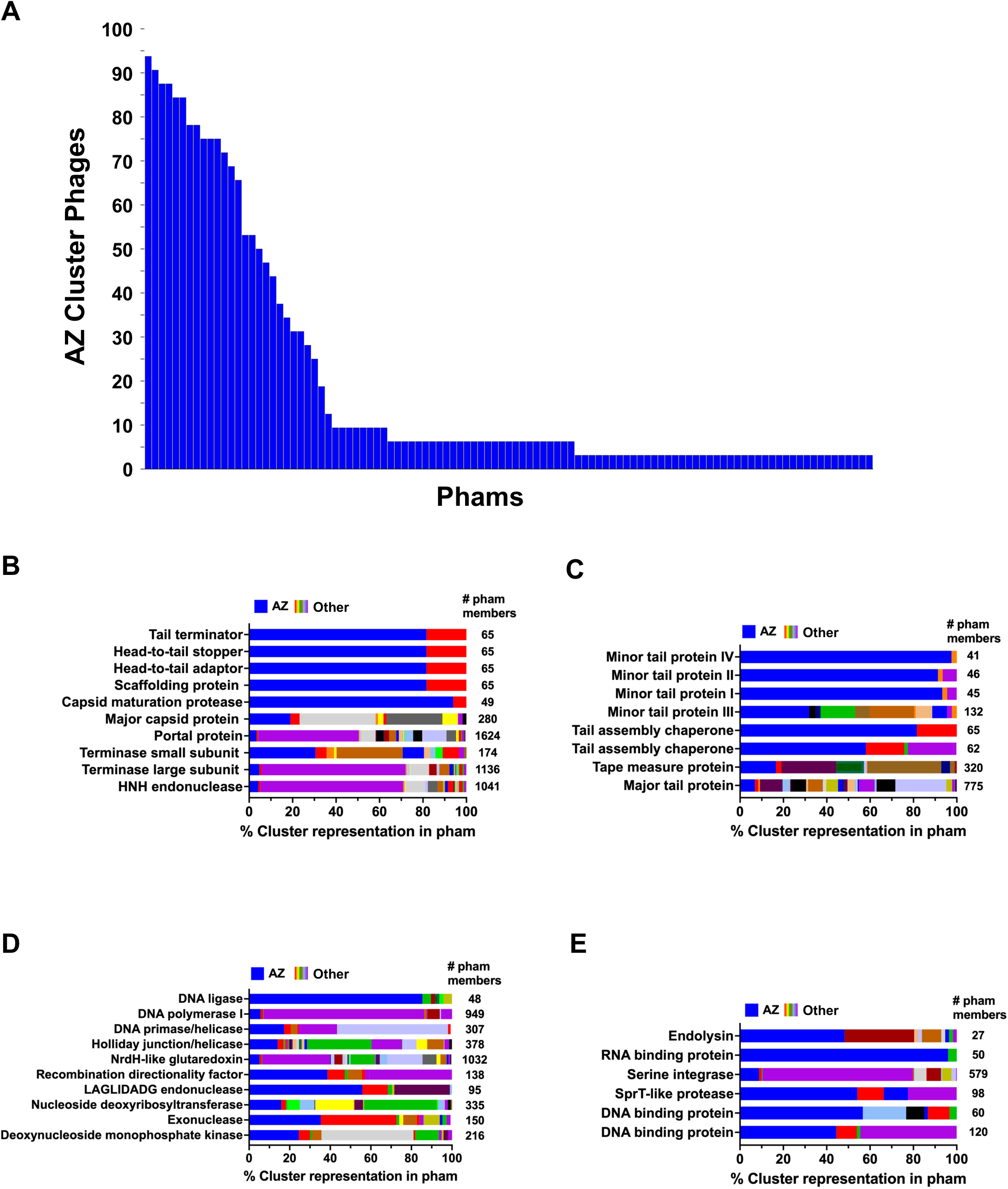
Representative Cluster AZ pham analysis. Pham analyses were performed using the Actinobacteriophage_4491 database. (**A**) Phams uniquely found in phages from 32 AZ cluster phages (n=105). Each bar represents a pham currently found only in AZ phages, with the percentage of AZ phages with each particular pham indicated on the Y axis. All unique phams code for proteins of unknown function. Figure generated with custom Javascript notebook (https://observablehq.com/@phage/cluster-specific-phamilies) using database Actinobacteriophage_4491. (**B-E**) Cluster distribution of selected phams with known functions, using phage Lizalica as an example (Fig. 2). Each row represents a pham (denoted here with specific functions), and each color represents the proportion of phages from various clusters that contain that pham. Cluster AZ is shown in blue; other clusters are indicated by a variety of colors. (B) Structural phams associated with capsid. (C) Structural phams associated with the tail. (D) Phams with DNA metabolism functions. (E) Phams in Integrase region with potential roles in lysogeny.

We analyzed the cluster distribution of phams in AZ genomes with assigned functions (**Fig. 10B-E**), using cluster AZ1 phage Lizalica as the comparator genome (**Fig. 2**). The 34 distinct phams in Lizalica illustrate the gene mosaicism in this cluster and reveal two different distribution patterns. One group of phams include a large number of phage members (7/34 phams had over 500 members; 4 of these had over 1000 members) and are widely distributed in a diverse set of actinobacteriophages, including those infecting other non-*Arthrobacter* hosts (**Figure 10B-E**; **Supplemental Table 3**). In some of these larger phams, there are representatives from as many as 30 different clusters, representing seven different genera of Actinobacterial hosts (**Supplemental Table 3)**. These large and widely distributed phams include every major functional category but are particularly represented in phams involved in DNA metabolism, many structural proteins and the serine integrase. These findings reveal several phams that have been widely conserved as Actinobacteriophages diverged into different clades and evolved to infect a diversity of hosts.

A second group of phams in AZ phages have fewer than 70 members, with a very narrow distribution, consisting of Clusters AZ, EH, and FP (**Fig. 10B-E; Supplemental Table 2**). These phams include minor tail proteins, a head-to-tail adaptor, a head-to-tail stopper, and a tail terminator. In addition, with the exception of the integrase, phams with functions potentially involved in gene regulation and lysogeny also have a narrow distribution and are mostly represented in the AZ cluster **(Fig. 10E)**. These findings highlight functions that may delineate individual clades and adaptation to hosts.

## Concluding remarks

Here we have described a comparative genomic analysis of a large population of Cluster AZ bacteriophages infecting *Arthrobacter* bacterial hosts, extending an initial study on a smaller group of phages (20). These phages share enough nucleotide similarity to be grouped into the same cluster, AZ, although there is a spectrum of gene content diversity that warrants dividing these phages into four subclusters. Since the analyses in this paper were completed, additional Cluster AZ phages have been isolated on Arthrobacter hosts as well as a new genus, *Curtobacterium* (https://phagesDB.org). This host genus is a member of the family Microbacteriaceae, to which the genus *Microbacterium* also belongs. Future studies of these Cluster AZ phages isolated on *Curtobacterium* may further support the relationship observed between Cluster AZ phages isolated on Arthroba*cter* and the Cluster EH *Microbacterium* phages, which are still the most closely related phages outside of Cluster AZ. This finding suggests that Cluster AZ and EH phages share an evolutionary history that is distinct from the rest of the *Arthrobacter* and *Microbacterium* phages. Although this relationship might suggest an expanded host range, we have not found that these phages can infect each other’s hosts (20) (Rudner, unpublished data). The continuum of diversity described in Cluster AZ phages is similar to that seen with the *Mycobacterium* phages, and our observations at the cluster and subcluster levels parallel those observed among *Mycobacterium* phages as additional phages were sequenced and analyzed (16,19,40). Since these studies were completed, many more *Arthrobacter* phages have been isolated, sequenced, and annotated, including many in cluster AZ (48). Further genomic analyses of these new phages will continue to provide new insights into phage biology and evolution.

## Materials and methods

### Bacterial strains and media

Phages were isolated on *Arthrobacter globiformis* (NRRL B-2979 or B-2880), *Arthrobacter atrocyaneus* (NRRL B-288), or *Arthrobacter sulfureus* (NRRL B-14730) hosts. Bacteria were cultured in PYCa media (1.0g yeast extract, 15g peptone, 2.5 mL 40% dextrose, and 4.5 ml 1M CaCl_2_ per L volume) at 30°C with constant shaking as previously described.

### Isolation of *Arthrobacter* phages

Phages were isolated directly or with enrichment from soil samples (https://seaphagesphagediscoveryguide.helpdocsonline.com/home). For direct isolations soil samples were shaken vigorously in phage buffer (10mM Tris-HCL, pH 7.5; 10mM MgSO_4_; 68.5mM NaCl; 1mM CaCl_2_) or PYCa media for 2 hours at 30°C, centrifuged to pellet the soil and debris, and the phage-containing supernatant was filtered with 0.22μm - 0.45μm filters to generate a lysate. Isolated plaques were visualized on double agar overlays containing PYCa 0.35-0.70% agar and *Arthrobacter* host and incubated at 30°C for 16-48 hours. For enriched isolations, filtered soil supernatant was prepared as above and incubated at 30°C with *Arthrobacter* host in PYCa for 1-5 days, refiltered with 0.22μm - 0.45μm filters and tested for the presence of phage. VroomVroom and Emotion were found by using mixed-enriched isolations in which a mixture of *A. sulfureus*, *A. atrocyaneus* and *A. globiformis* B-2880 and B-2979 were added to the filtered soil supernatant. For both enriched soil samples and the direct soil samples, phages were purified twice through at least two rounds of plaque purification and high titer lysates were generated using methods described previously for Mycobacterial hosts (27,49). These high titer lysates were archived at -80°C and used for all further analyses.

### Transmission electron microscopy

High-titer lysate of Crewmate was stained with 1% uranyl acetate on a Carbon Type-B, 300 mesh, copper grid (Ted Pella #01813). The grid was visualized at the uOttawa TEM core facility at 120kV on a JEOL JEM-1400plus TEM. Scale bars were added and the image was adjusted using Fiji (ImageJ). Additional TEM images of the AZ phages can be found on PhagesDB (https://phagesdb.org).

### DNA extraction, genome sequencing and genome annotation

Phage DNA was extracted using Promega kit A1120 according to the manufacturer’s instructions. Phage genomes were sequenced at the University of Pittsburgh phage genomics facility using Illumina platforms to at least 20-fold coverage. Shotgun reads were assembled *de novo* with Newbler versions 2.1 to 2.9. Assemblies were checked for low coverage and discrepant areas and targeted Sanger reads were used to resolve weak areas and identify genome ends. All genome sequences are publicly available at phagesdb.org and in GenBank (Table 1).

Genomes were annotated using DNA Master (http://cobamide2.bio.pitt.edu/) and the Phage Evidence Collection And Annotation Network (PECAAN; https://discover.kbrinsgd.org/), (which incorporate data from GLIMMER (50), GeneMark (51), Starterator (52), BLAST (53), HHPred (54), SOSUI (55), TMHMM (56), DeepTMHMM (57), and Phamerator.org (33), using database Actinobacteriophage_4491. Aragorn (58) and tRNA-ScanSE (59) were used to identify tRNA genes.

### Harmonization of annotated phages

Phage annotations were “harmonized” to ensure consistency in gene starts and function assignments across the cluster and resolve any discrepancies that arise due to the large number of annotators involved and the dynamic nature of the data. Harmonization was performed manually by creating a spreadsheet comparing all start sites and functions for each pham and then using an Observable script that automatically creates this spreadsheet, and identifies all tandem start sites and gaps between genes that may harbor missing ORFs (**Fig. S1**; https://observablehq.com/@phage/pharmonizer).

### Comparative genomic analyses

To create the gene content network phylogeny, pham data was acquired from the database Actinobacteriophage_4991 and analyzed using Phamnexus (http://github.com/chg60/phamnexus) and visualized using SplitsTree (60,61). Gene content similarity (GCS) analyses were conducted by using pdm_utils (https://academic.oup.com/bioinformatics/article/37/16/2464/5998668) to export pham data from the Actinobacteriophage_4991 database. Prism v10 (GraphPad, Boston, MA, USA) was used to visualize GCS data and perform statistical analyses.

### Multiple sequence alignments

To evaluate relationships between minor tail protein sequences, the minor tail proteins were aligned using MUSCLE on the EMBL-EBI website (62). The alignment was imported into iTOL (https://itol.embl.de/about.cgi) (63) to create the unrooted tree diagram. To compare the sequences of the endolysin genes, the protein sequences for the genes were downloaded as a multifasta file from PhagesDB. A multiple sequence alignment was generated using MUSCLE in Unipro UGENE v48.0 (64). The region of the MSA containing the proline-alanine repeat was noted as a region of interest.

### Alphafold3 structure predictions

Alphafold3 was run on the Alphafold server (65). The model_0.cif was searched on the FoldSeek server (44). Structures were compared and colored on Pymol v2.5.4 (66).

## Supplemental figure legends

**S1 Fig. Automating the organization and analysis of cluster data using Pharmonizer**. Harmonization uses Pharmonizer to access data from Phamerator and analyzes genes for tandem start sites, and discordant protein functional assignments, identifies large gaps to be re-investigated and compares pham gene lengths to flag potential start site discordance. Pharmonizer can output DNA sequences for bioinformatic analysis and creates an organization grid that allows a simplified comparison of each pham and tracking of changes to be made in GenBank files.

**S2 Fig. BaileyBlu genome map.**

Genome of Cluster FP phage BaileyBlu, a relative of Cluster AZ phages. Genes are depicted as boxes and are colored according to their phamily designations (phams). White boxes represent genes with no close relatives in the database.

**S3 Fig. Genomic differences between phages Maureen and Liebe.**

**A)** NCBI BlastN comparison of Liebe and Maureen reveals four nucleotide differences. **B)** Liebe *19, 40* and *55* show single nucleotide polymorphisms that change a single amino acid in each protein. Liebe *49* deletes a single nucleotide, resulting in a change of start site in the gene and a truncated protein. **B)** In Liebe, a cytosine insertion at bp 36,432 causes a frameshift if the ATG start is used. A truncated gp49 protein of 163aa can be produced if the GTG start is used at bp 36,477.

**Suppl Table 1 (GCS numerical values)**

**Suppl Table 2 (AZ Endolysin Types)**

**Suppl Table 3 (Pham Analyses)**

## Acknowledgments

We thank Debbie Jacobs-Sera and Graham Hatfull for their helpful comments on the manuscript and Christian Gauthier for the construction of the Actinobacteriophage_4491 database. We also thank Daniel Russell and Rebecca Garlena for phage sequencing and data curation. We are grateful for programmatic assistance from Billy Biederman, Danielle Heller, Vinesh Sivanathan and David Asai, and to the SEA-PHAGES and PHIRE programs for programmatic support. We especially thank all the students at our institutions who participated in the isolation of these unique phages. K.P.F was funded by an OISB/NRC graduate fellowship and a Canada Graduate Scholarship award, Z.M. was funded by an Ontario Graduate Scholarship, A.D.R. was supported by an NSERC Discovery Grant.

