## Supplementary figures and images for "Comparative genomic analysis of Cluster AZ *Arthrobacter* phages"

### Supplemental Figure S1

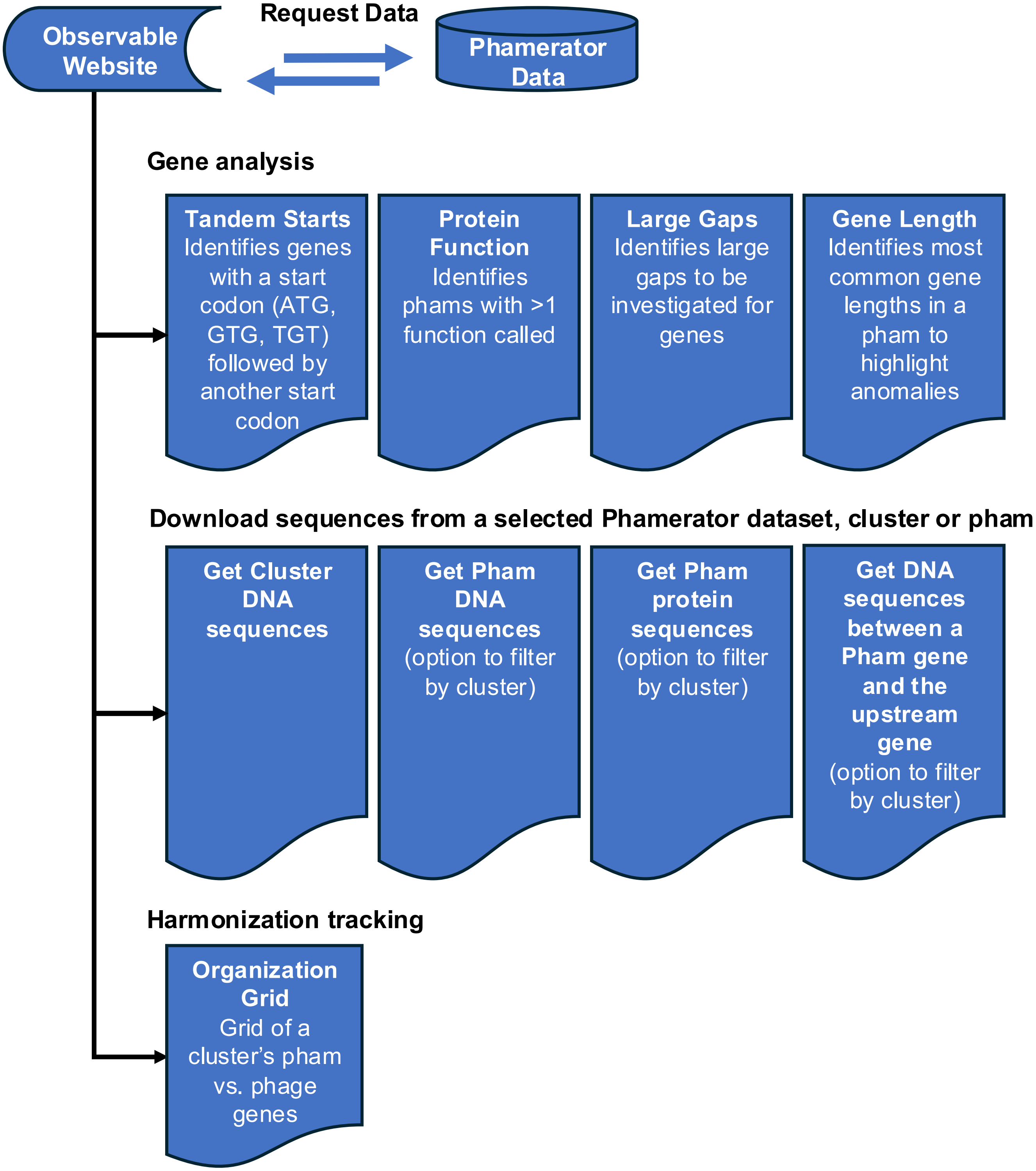

### Supplemental Figure S2

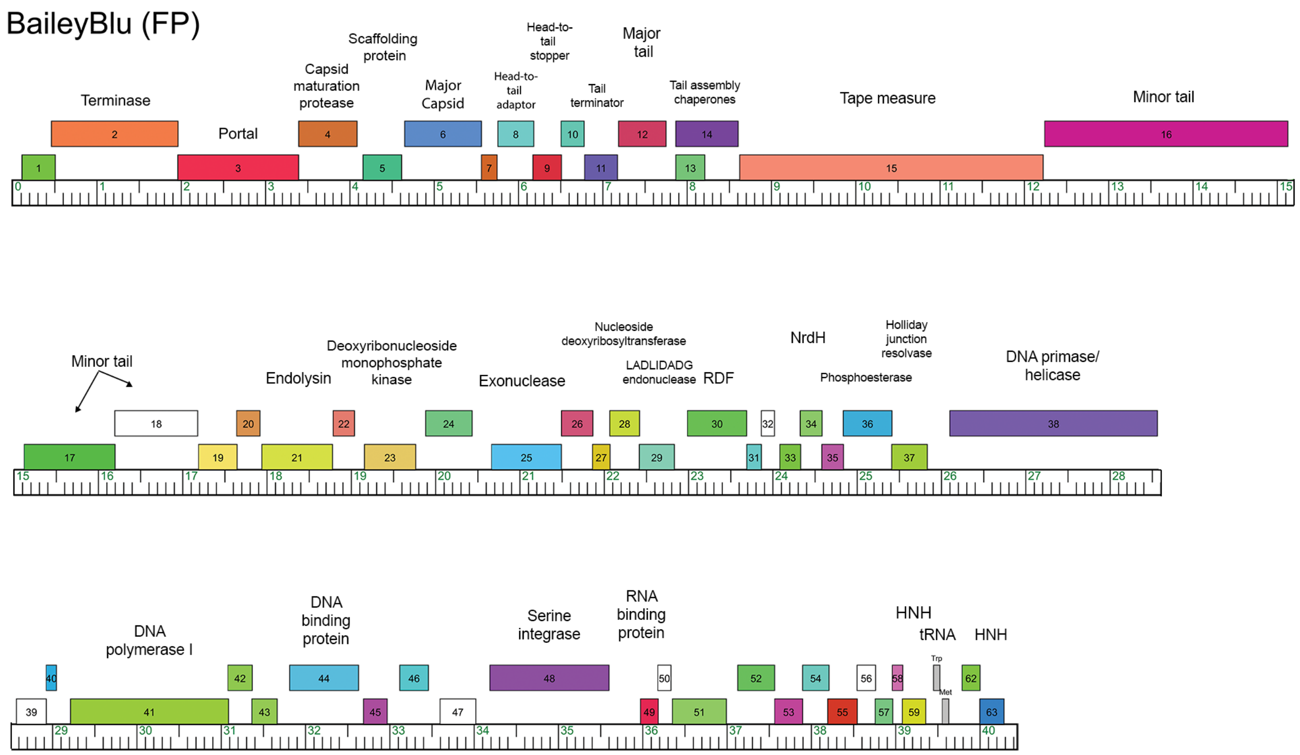
