## Supplemental Figure S3 for "Comparative genomic analysis of Cluster AZ *Arthrobacter* phages"

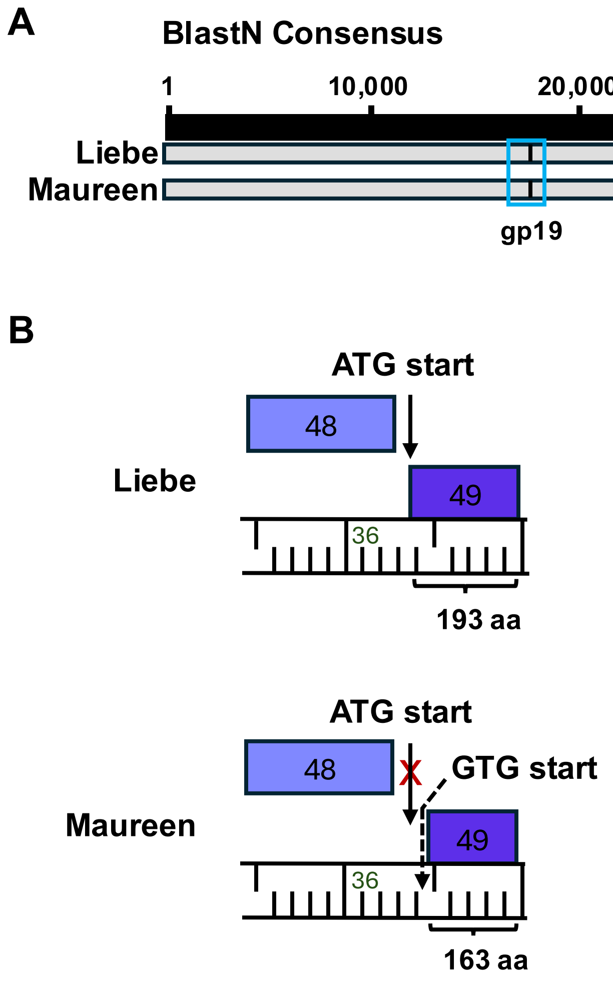


C

| Maureen Genome vs Variants in Liebe | | | | | |
|---|---|---|---|---|---|
| Gene (gp) | Function | Sequence | | Protein | |
| | | Locus | Nucleotide change | Amino acid change | Length (aa) |
| 19 | NKF | 17,524 | A>G | G > D | 1,076 |
| 40 | NKF | 30,384 | T>G | G > C | 257 |
| 49 | SprT-like protease | 36,432 | delC | p.(Met1Valext) | 196 (Maureen)<br>163 (Liebe) |
| 55 | NKF | 40,104 | G>A | T > A | 311 |
